# Temporal Parameters Determine the Efficacy of Vagus Nerve Stimulation Directed Neural Plasticity

**DOI:** 10.1101/2024.12.16.628694

**Authors:** Juliet J. A. Addo, Connor L. Neifert, Tanya T. Danaphongse, Stephanie T. Abe, Vikram Ezhil, Michael P. Kilgard, Seth A. Hays

## Abstract

Combining vagus nerve stimulation (VNS) with rehabilitation represents an emerging treatment for a range of neurological disorders, and identifying stimulation parameters that maximize the effects of VNS may provide a means to optimize this therapy. A number of prior studies show that varying the intensity of stimulation, which influences the amount of neuromodulatory activation in response to VNS, determines the strength of VNS-dependent enhancement of synaptic plasticity in cortical circuits. The impact of the temporal parameters of stimulation, such as the frequency and distribution of pulses within a stimulation train, remains underexplored. In this study, we evaluated how varying these temporal parameters impacts the magnitude of VNS-directed plasticity. In the first experiment, rats received trains of VNS at one of three moderate pulse frequencies (20, 30, or 45Hz) timed to occur with movement during training on a simple motor task. After five days of training, we evaluated the cortical movement representations using intracortical microstimulation. All three moderate pulse frequencies produced equivalent increases in the cortical representation of the paired movement compared to sham stimulation. In a second experiment, we used a similar paradigm to explore whether burst stimulation would enhance VNS-dependent plasticity. Unexpectedly, we found that both burst stimulation or a matched number of pulses distributed evenly in time failed to produce significant enhancement of plasticity compared to sham stimulation, whereas moderate pulse frequency stimulation did. These findings illustrate the importance of the temporal dynamics of stimulation in determining the effects of VNS and provide guidelines for designing novel VNS sequences.

## Introduction

Vagus nerve stimulation (VNS) combined with rehabilitation has emerged as a promising treatment for a range of neurological disorders, including stroke and spinal cord injury^1–12^. This approach is premised on VNS-dependent activation of neuromodulatory circuits in order to facilitate synaptic plasticity in networks engaged by the concurrent rehabilitation^8,13^. The magnitude of neuromodulatory activation is dependent on the electrical stimulation parameters of VNS^14,15^. Consequently, selecting VNS parameters that optimize the degree of activation holds promise to increase the efficacy of VNS therapy.

The vast majority of studies have, and reasonably so, focused on changing the intensity of stimulation to alter the degree of VNS-driven neuromodulator release. Indeed, the intensity of stimulation directly influences the magnitude of noradrenergic locus coeruleus (LC) activation^14^. Paradoxically, one of the primary findings from these studies is that stimulation intensity exhibits an inverted-U relationship with VNS-dependent plasticity. As such, moderate intensity stimulation enhances plasticity, whereas both lower and higher intensity stimulation fail to do so. Moreover, this inverted-U relationship holds true for the actions of VNS on recovery in a number of disease models^7,14,16–22^. Collectively, this leads to two conclusions: First, moderate intensity stimulation appears to be the most effective. Second, the inverted-U relationship complicates parameter selection, such that more activation does not strictly produce more plasticity and recovery. As a result, a more comprehensive evaluation of other stimulation parameters beyond intensity is justified.

The temporal parameters of stimulation, which are also known to predictably influence LC activation and would therefore be expected to influence plasticity^14^, remain largely unexplored. Initial evidence using paired VNS points to the importance of temporal parameters, but the wide range of parameter variation in these studies provides little insight into the contours of the relationship between pulse frequency and plasticity and may have missed optimally effective moderate pulse frequencies^23,24^. As a practical matter, varying temporal stimulation parameters by using different train durations, pulse frequencies, and burst patterns, has been useful in identifying more effective stimulation regimens for other neuromodulation therapies, such as deep brain stimulation and spinal cord stimulation^25–27^. Similar optimization of paired VNS therapy may be possible by exploring alternative temporal paradigms of stimulation.

In this study, we performed two experiments to evaluate different aspects of temporal parameters on VNS-dependent plasticity. In the first study, we characterized the effect of varying pulse frequency on the degree of VNS-dependent plasticity. We sought to determine if pulse frequencies nearby, but distinct from, the commonly used 30Hz would be more or less effective. In a second study, we explored whether varying the train duration and bursting of pulses would influence the degree of VNS-dependent plasticity. Our findings reiterate the importance of considering temporal parameters when utilizing VNS and provide a basis for selecting stimulation paradigms to improve efficacy.

## Methods

Experimental procedures, statistical comparisons, and exclusion criteria were preregistered prior to data collection commenced (https://osf.io/rxnm7, https://osf.io/ydm3w).

### Animals

Ninety-one adult female Sprague Dawley rats aged approximately 6 months old, weighing between 250g to 300g, were obtained from Charles River Laboratories and used in this study. All animals were housed in a reversed 12:12 hour light-dark cycle. Rats were food restricted on weekdays during shaping and behavioral training with ad libitum access to food on weekends. Throughout the study, all rats were kept at or above 85% of their initial body weight upon commencement of behavioral testing.

### Behavioral training

Rats were trained on a simple automated behavioral task that enabled VNS to be paired with chewing^21^. During each session, rats were placed into an acrylic cage outfitted with a food pellet dispenser with a nose poke food dispenser. Rats were trained to nose poke to receive a reward food pellet (45mg dustless precision pellet, Bioserv, Frenchtown, NJ). At the start of a session, a food pellet was delivered to the food dispenser^21^. Nose poke events were detected when rats broke an infrared beam positioned in the food dispenser to retrieve the pellet. Once the infrared beam was intercepted, another pellet was dispensed after an 8 second delay^21^. This continued until 1 hour elapsed or 100 food pellets were dispensed^21^. Training sessions were run twice daily for 5 days per week with at least a 2-hour break between each daily session until the rats reliably performed the task^21^. Rats were given 10g supplemental food pellets if they did not receive at least 100 food pellets in a day during behavioral training sessions to maintain their weight.

### Vagus nerve stimulator cuff implantation

Once proficient at the task, rats underwent surgical implantation of a custom platinum-iridium bipolar stimulating cuff electrode on the left cervical vagus nerve and headmount as described in previous studies^1–3,5–7,10,12–14,20,21,23,28,29^. All surgeries were conducted using aseptic techniques under general anesthesia. Rats were anesthetized using a cocktail of ketamine hydrochloride (50mg/kg, i.p.), xylazine (20mg/kg, i.p.), and acepromazine (5mg/kg, i.p.), and anesthesia levels were stabilized throughout the surgical procedure by a combination of the assessment of breathing rate, toe pinch reflex, whisking, and the use of a pulse oximeter to track heart rate and blood oxygen saturation^20,21^. Once anesthetized, rats were placed in a stereotaxic frame, and an incision was made along the midline of the head to expose the skull. Four bone screws were inserted into the skull at points along the lambdoidal and sagittal sutures to serve as an anchor for the 2-channel connector headmount. The headmount was anchored to the screws and skull using acrylic. The rat was then removed from the stereotaxic frame and placed in supine position. A 10mm incision was made parallel to the dorsal midline, centered approximately 5mm lateral and 5mm anterior to the top of the rat’s sternum. The underlying musculature was blunt dissected to expose the left cervical vagus nerve. The vagus nerve was separated from the carotid sheath and placed in the cuff. The cuff leads were then tunneled subcutaneously dorsally along the neck and inserted to the 2-channel connector headmount.

Electrode functionality was confirmed with observation of a ≥5% drop in the blood oxygen saturation in response to a 0.5s stimulation train of VNS consisting of 0.8mA, 100μs biphasic pulses at 30Hz, as described in previous studies^20,21,30^. After confirming cuff functionality, all incisions were sutured closed, and rats were given 4mL subcutaneous injections of 50:50 0.9% saline 5% dextrose solution. All rats underwent a 7-day recovery period before returning to behavioral training^20,21^.

### Vagus Nerve Stimulation

After recovery, rats were randomly assigned to either the Sham (no stimulation; n = 10), 20Hz (n=10), 30Hz (Standard VNS) (n=10), 45Hz (n=10), Burst VNS (n=10), or Dispersed VNS (n=10) group. Training timelines for individual rats were interleaved such that data collection for experiments 1 and 2 happened simultaneously. During behavioral task sessions, all rats were connected to a stimulation cable via the headmount. In the initial sessions following implantation, rats were allowed to habituate to the stimulation cable until they performed 200 successful trials per day. Once acclimated, rats underwent twice daily sessions of behavioral training for 5 days, during which they received VNS or sham stimulation based on their group assignment (Fig. 1). Rats receiving sham stimulation were connected to the cable, but did not receive stimulation. For experiment 1, each 0.5s stimulation train consisted of 100μs biphasic pulses delivered at 0.8mA at a frequency of either 20Hz, 30Hz or 45Hz. For experiment two, the number of pulses were conserved for the Dispersed and Burst VNS groups: Dispersed VNS was delivered at a frequency of 7.5Hz over a duration of 2s, and Burst VNS delivered 125ms bursts of 30Hz stimulation separated by 375ms pauses over a duration of 2000ms. For all groups, VNS was delivered 3s after the infra-red beam was intercepted by a rat, ensuring that stimulations were delivered concurrent to chewing, as seen in previous studies^20,21^. A digital oscilloscope (PicoScope 2204A, PP906, Pico Technology) was used to assess the voltage delivered across the vagus nerve during each stimulation to ensure cuff functionality. All data for the total amount of VNS delivered and intervals between stimulations is available as supplementary material (Table S1).

**Figure 1.**
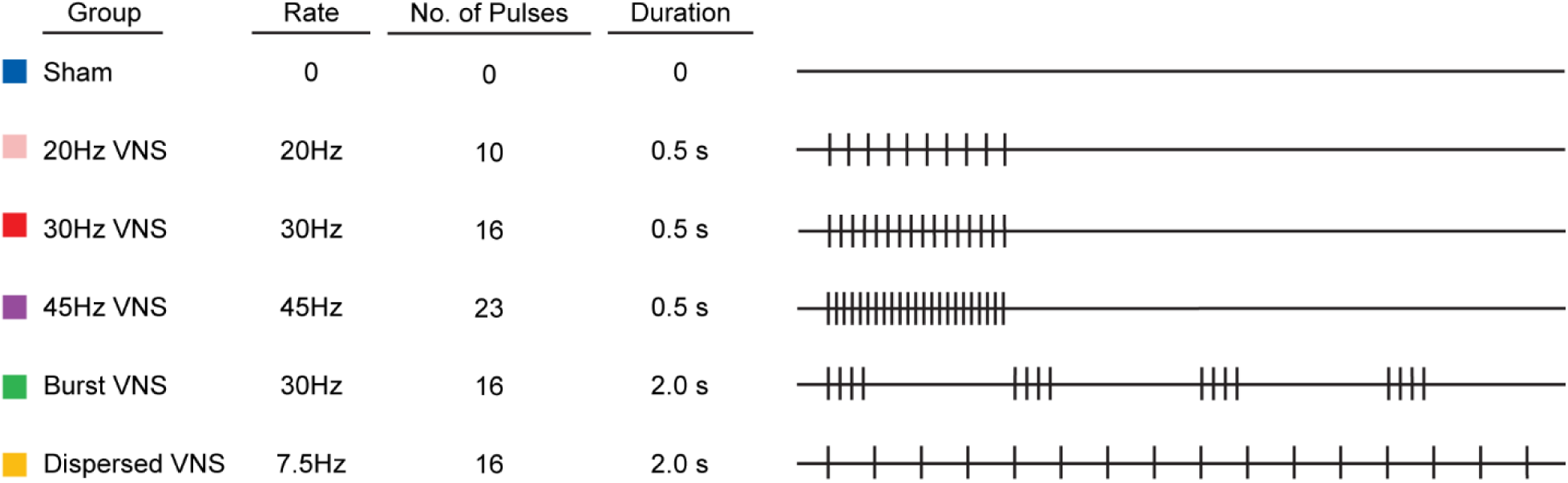
Schematic description of the temporal parameters of VNS paradigms tested in this study. Each group is depicted as a row. The right panel illustrates a single train of VNS for each group. Vertical ticks represent individual pulses within a train.

### Intracortical microstimulation (ICMS) mapping

Approximately 24 hours after the final session of behavioral training with VNS or sham stimulation, rats underwent ICMS to derive cortical movement representations. Prior to the start of ICMS, nerve cuff functionality and nerve activation were assessed as described earlier, irrespective of their group.

ICMS was used to produce functional representative maps of the left motor cortical area according to established protocols^19,29,31,32^. Rats were first anesthetized using an intraperitoneal injection of ketamine hydrochloride (75mg/kg) and xylazine (5mg/kg). Supplemental doses of ketamine (25mg/kg) and xylazine (1.5mg/kg) were administered as necessary throughout the procedure to maintain a consistent rate of anesthesia, informed by pulse-oximetry observations (Smith’s Medical, SurgiVet) of the rat’s blood oxygen saturation and heart rate, and observations of toe pinch reflex and vibrissa whisking ^20,21^. After initial anesthesia, rats were placed in a stereotactic apparatus. A craniotomy and durotomy exposed the left motor cortex (4mm to −3mm AP and –0.25mm to –5mm ML from bregma)^20,21^. A small incision was also made in the cisterna magna to prevent cortical swelling.

A tungsten electrode with an impedance of approximately 0.7MΩ (UEWMEGSEBN3M, FHC, Bowdoin, ME) was lowered to a depth of 1.8mm into a pre-selected site in the left motor cortex with the aim of targeting motor outputs in layer V. Stimulation sites were then chosen at random on a grid with sites set 500μm apart from each other. The subsequent stimulation sites were placed at least 1 mm away from the previous site. Stimulation consisted of a 40ms pulse train of 10 monophasic 200μs cathodal pulses. Stimulation started at 10μA and was increased by 10μA until a movement was observed or until a maximum of 250μA was reached. ICMS was conducted by two experimenters who were blinded to the experimental groups, as previously described^19–21,28^. The first experimenter was responsible for placing the electrode and recording the data for each site. The second experimenter, blinded to the electrode placement delivered stimulations and categorized observed movements as either jaw, neck, vibrissa, forelimb, or hindlimb. The primary outcome of this study was the area of the motor cortex producing jaw movements. All other movement representations were assessed as secondary outcome measures. All data from ICMS is available as supplementary material (Table S2, Fig. S1A-F).

### Subject exclusion

Exclusion criteria were preregistered prior to data collection. 60 subjects were assessed in the results of the study out of a total of 91 subjects. Of the 31 subjects excluded from final analysis, 2 subjects were excluded due to a non-functional stimulating cuff identified by digital oscilloscope readings exceeding 40V peak-to-peak, 12 subjects were excluded due to mortality during surgical procedures, 10 subjects were excluded due to mechanical failure of the headmount during behavioral training, and 4 subjects were also excluded due to complications in VNS delivery. 3 subjects were excluded due to anesthesia complications during the map, defined by hourly observations of blood oxygen saturation, heart rate, and the strength of toe pinch reflex (rated from 0 (no response) to 6 (a rapid, awake-like large contraction)). Timepoints were averaged for each measure. Average blood oxygen saturation (%) was divided by 100, heart rate by 350bpm, and toe pinch strength was divided by 6. The three fractions were then summed, and a rat with final score of >2.5 was deemed overall too light on anesthesia to generate a reliable map.

### Statistics

Outcomes and planned comparisons were preregistered and defined a priori. A D’Agostino & Pearson test confirmed data normality. The primary outcome measure of this study was the area of the motor cortex eliciting jaw movements. All areas eliciting other movements were assessed as secondary outcome measures. Changes in area across our experimental groups were assessed using One-way ANOVAs and unpaired two-tailed t-tests to compare individual groups. Statistical tests for each comparison are noted in the text. Data is reported in the text and figures as mean ± standard error of the mean (SEM).

## Results

In the first experiment, we sought to evaluate the effect of pulse frequency on VNS-dependent plasticity in motor cortex. To do so, rats received training on a simple behavioral task during which VNS was administered during jaw movement at a fixed pulse frequency of 20Hz, 30Hz, or 45Hz (Fig. 1). After five days of training, cortical movement representations were derived with intracortical microstimulation.

Comparison of the stimulation groups to sham revealed a significant increase in cortical jaw representation with VNS (Fig. 2A; Unpaired t-test; t(38) = 3.4, p = 0.019). As seen in previous studies, the VNS-dependent enhancement of plasticity was specific to the paired movement. Other movement representations and the total map area were unchanged across groups (Fig. S2A,B; One-way ANOVA, total map area: F[3,36] = 0.7226, p = 0.5451; forelimb: F[3, 36] = 0.3138, p = 0.8153; p = 0.1093; vibrissa: F[3, 36] = 0.1405, p = 0.9351; neck: F[3, 36] = 1.2667, p = 0.3004; hindlimb: F[3, 36] = 0.1309, p = 0.9411). These findings demonstrate that moderate pulse frequencies of VNS produce comparable enhancements of cortical plasticity specific to the paired movement, consistent with prior studies^1,20–22,29^.

**Figure 2.**
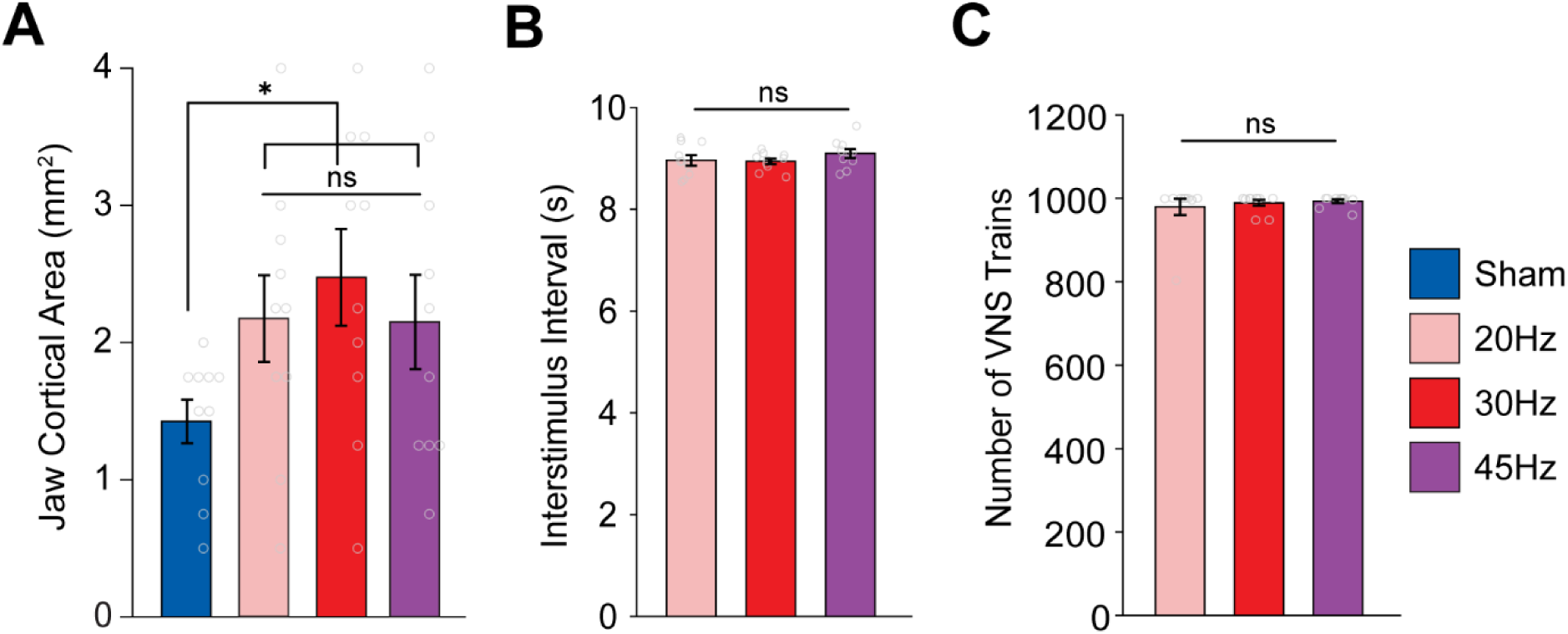
Moderate frequencies of VNS produce comparable enhancement of plasticity. (A) Moderate frequencies of VNS paired with chewing significantly increase the cortical representation of jaw muscles compared to Sham. (B, C) No differences in the interval between stimulation trains nor the total number of trains were observed between VNS groups. Bars represent mean ± SEM. Gray circles show individuals subjects. * denotes p < 0.0167; n.s. not significant.

We next sought to determine if differences in behavior or stimulation could account for the similar effects of VNS. We also observed no differences in the total amount of VNS delivered or in the interval between stimulations across the VNS groups (Fig 2B,C; One-way ANOVA, total amount of VNS: F[2,27] = 0.3197, p = 0.7291; time between stimulations: F[3, 36] = 0. 98027, p = 0. 3882). Additionally, the threshold to elicit movement during ICMS was similar across groups (Fig. S2C; One-way ANOVA, F[3, 36] = 0.4473, p = 0.7207). These findings confirm that the comparable enhancement of jaw representation across VNS groups can be ascribed to a similar magnitude of VNS effect across these moderate frequencies.

Because pulse frequency across the moderate range did not affect the magnitude of VNS-dependent plasticity, we next sought to determine whether burst stimulation would provide a means to generate greater enhancement of plasticity. In this experiment, rats received a similar pairing strategy, but VNS was delivered either in 4-pulse bursts at 30Hz over a 2 second train (Burst) or a matched number of pulses distributed evenly over a 2 second train (Dispersed). We observed a significant effect of group (Fig. 3A; One-way ANOVA: F[3,36] = 2.9211). Unexpectedly, burst stimulation failed to enhance plasticity compared to equivalent training without stimulation (Fig. 3A; Unpaired t-test Sham v. Burst; t(18) = 0.8956, p = 0.3823). The same number of pulses distributed over 2 seconds (i.e., at 7.5Hz), also failed to enhance plasticity (Unpaired t-test Sham v. Dispersed; t(18) = 1.3606, p = 0.1904). Total map area and other movement representations were unchanged across groups (Fig. S3A,B; One-way ANOVA, total map area: F [3,36] = 0. 9248, p = 0.4387; forelimb: F[3, 36] = 0. 1672, p = 0.9178; vibrissa: F[3, 36] = 1.2405, p = 0.3093; neck: F[3, 36] = 0.3373, p = 0.7985; hindlimb: F[3, 36] = 0.0969, p = 0.9613). No differences in ICMS threshold were observed between groups (Figure S3C) (One-way ANOVA, F [3,36] = 0.173, p = 0.9140). Additionally, no differences in the total amount of VNS delivered or in the time between stimulations were observed between groups (Fig. 3B,C; One-way ANOVA, total amount of VNS: F[2,27] = 0. 1815, p = 0. 8350; time between stimulations: F[3, 36] = 0. 5867, p = 0.5631). Given that the number of VNS pulses in a train for the Burst and Dispersed groups is matched to the 30Hz VNS group yet they do not change cortical representations, these findings highlight the impact of the temporal dynamics of pulses within a stimulation train on VNS-dependent plasticity.

**Figure 3.**
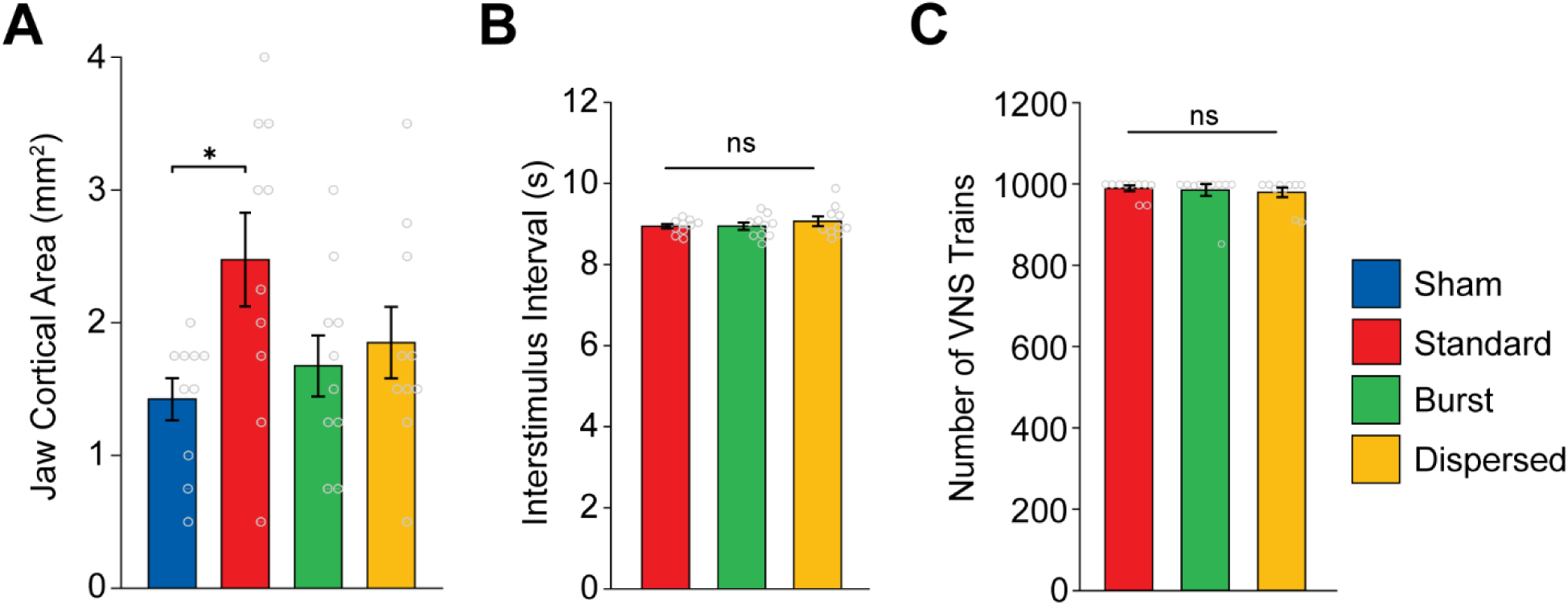
Temporal dynamics of pulses within a train influence the magnitude of VNS-dependent plasticity. (A) Standard (30Hz) VNS paired with chewing significantly increases the cortical representation of jaw musculature compared to Sham. Short bursts of VNS bursts within a train (Burst) and the same number of pulses evenly distributed across the train (Dispersed) did not increase cortical area compared to Sham. (B, C) No differences in the interval between stimulation trains nor the total number of trains were observed between VNS groups. Bars represent mean ± SEM. Gray circles show individuals subjects. * denotes p < 0.0167; n.s. not significant.

## Discussion

VNS combined with various forms of training represents a means to improve recovery in individuals with neurological disorders, and optimizing the stimulation parameters to maximize efficacy is of key importance^16^. In this study, we sought to determine if the temporal parameters of stimulation would influence the degree of VNS-dependent enhancement of plasticity. We found that moderate pulse frequencies, across an approximate 2-fold range, appear equivalently effective. Additionally, we find that burst or distributed stimulation are less effective than moderate pulse frequencies. These findings highlight the importance of the temporal parameters of stimulation on the effectiveness of VNS.

In a first experiment, we explored whether pulse frequencies (i.e., the rate at which pulses are delivered during a stimulation train) could increase the magnitude of VNS-dependent plasticity. Prior studies indicate that, compared to the often-used 30Hz, much lower (7.5Hz) or much higher (120Hz) pulse frequencies failed to promote VNS-dependent plasticity^23,24^. This mirrors the inverted-U relationship between VNS-dependent plasticity and stimulation intensity that has been widely observed, in which moderate parameters produce greater efficacy than lower or higher parameters^16^. While these relatively extreme conditions were not effective, we reasoned that perhaps there was a local maximum that may be observed with pulse frequencies near the effective and broadly used 30Hz pulse frequency. We found that 20Hz and 45Hz produced an enhancement of plasticity comparable to 30Hz. While we cannot exclusively rule out the possibility that other pulse frequencies may be more effective, the similarity across these parameters indicates that there is not an obvious range of more effective frequencies.

An additional motivating premise for this experiment was to probe the range of pulse frequencies over which VNS would promote enhanced plasticity. Evidence from a number of studies shows that there is a relatively narrow range of effective stimulation intensities^7,17,18,20,21^. Dense sampling demonstrates that while 0.8mA stimulation enhances plasticity, 0.6mA and 1.0mA are less effective, highlighting the restricted range of effective intensities^21^. Our findings in this study indicate that 20Hz and 45Hz produce a similar enhancement of plasticity as 30Hz, indicating a broader range of effective pulse frequencies compared to intensities. Collectively, these findings demonstrate that across an approximately 2-fold range of moderate pulse frequencies, VNS combined with training produces a similar magnitude of plasticity enhancement and likely indicates that optimizing efficacy of VNS should focus on other stimulation parameters.

Because pulse frequency did not provide a clear means to optimize efficacy, in a second experiment, we explored whether changing the dynamics of pulses within a train would influence the magnitude of VNS-dependent plasticity (Fig. 1). Burst patterns, which are generally based on short epochs of pulses interspersed with silent intervals, have found favor for some forms of neuromodulation, and prior studies show that high frequency VNS may be effective. We found that neither burst stimulation nor a matched amount of stimulation distributed over the same train duration enhanced plasticity compared to sham stimulation. These findings illustrate two primary points. First, because all groups received a matched number of pulses at a fixed intensity and differed only in the temporal distribution of these pulses, we conclude that the temporal dynamics of the pulses within a VNS train determine its efficacy. These findings add to the existing corpus of data that the amount of total charge delivery, as defined by stimulation intensity and number of pulses, influences the magnitude of VNS-dependent plasticity and reiterates the importance of considering temporal patterns when selecting stimulation trains. Second, our results indicate that burst stimulation, at least within the constraints of parameters investigated in this study, are unlikely to produce greater clinical efficacy for paired VNS therapy.

VNS therapy is premised on the concept that stimulation drives rapid, phasic activity in the locus coeruleus, which releases norepinephrine to facilitate plasticity in networks activated during training^33–35^. The stimulation parameters dictate the magnitude and timing of locus coeruleus activation, which is reflected in the impact these parameters have on VNS-dependent plasticity^14^. Mirroring the inverted-U relationship between stimulation intensity and plasticity, our results are consistent with the notion that moderate pulse frequencies effectively enhance plasticity, while low and high pulse frequencies do not^23,24^. Collectively, these findings highlight that the dynamics, not just the total amount, of neuromodulatory activity in response to VNS influences plasticity. The engagement of noradrenergic receptors, in particular alpha-adrenergic receptors, is central to VNS-dependent enhancement of plasticity^35^. The magnitude and dynamics of stimulation, which determine the profile of neuromodulator release, thus likely determine which, when, and how many receptors are activated.

Combining VNS with rehabilitation has emerged as a treatment strategy for a number of disorders, culminating with FDA approval for the treatment of chronic stroke in 2021^8,36^. Identifying stimulation paradigms that maximize plasticity represents a means to potentially optimize the clinical benefits of this approach. The present study provides rationale and guidelines for selecting stimulation frequencies, and more broadly, reinforces the importance of considering the temporal profile when optimizing stimulation paradigms.

## Author Contributions

**J.J.A.A. and C.L.N.**: Equal contribution to Conceptualization, Methodology, Investigation, Analysis, Writing – original draft. **T.T.D.:** Investigation, Resources. **S.T.A.:** Investigation. **V.E.:** Investigation. **M.P.K.:** Conceptualization, Methodology. **S.A.H.:** Conceptualization, Methodology, Funding acquisition, Writing – review & editing, Project administration.

## Acknowledgements

We would like to thank Alfonso Reyes for the construction of VNS stimulator cuffs. We also thank Joseph Montefalcon, Swathi Pillai, Nanditha Niranjan, Ayesha Yousaf, Aadil Razack, Vishal Kumar, Praniya Jakkamsetti, Armaan Somaney, Roxanne Sanchez, and Vismaya Joseph for assisting with behavioral training and animal care. We would like to thank Robert Morrison for guidance on experimental methods, and technical expertise. We would also like to thank David Pruitt for developing behavioral programs and for guidance with analytical methods.

