## Supplemental Information for "Temporal Parameters Determine the Efficacy of Vagus Nerve Stimulation Directed Neural Plasticity"

**Supplementary Material:**

Table S1 – VNS delivery data for all subjects

Table S2 –ICMS data for all subjects

Figure S1 – Raw ICMS maps for all subjects

(A) Sham
(B) 20 Hz VNS
(C) 30 Hz Standard VNS
(D) 45 Hz VNS
(E) Burst VNS
(F) Dispersed VNS

Figure S2 – Other movement representations and ICMS thresholds are not changed by moderate VNS frequencies

Figure S3 – Other movement representations and ICMS thresholds are not changed by burst and dispersed VNS paradigms

### Table S1. VNS delivery data for all subjects

| Subject | Group | Average Interstimulation Interval (s) | Total Number of Stimulations |
|---|---|---|---|
| **S1** | Sham | n/a | n/a |
| **S2** | Sham | n/a | n/a |
| **S3** | Sham | n/a | n/a |
| **S4** | Sham | n/a | n/a |
| **S5** | Sham | n/a | n/a |
| **S6** | Sham | n/a | n/a |
| **S7** | Sham | n/a | n/a |
| **S8** | Sham | n/a | n/a |
| **S9** | Sham | n/a | n/a |
| **S10** | Sham | n/a | n/a |
| **S11** | 20 Hz VNS | 8.943 | 1000 |
| **S12** | 20 Hz VNS | 8.548 | 1000 |
| **S13** | 20 Hz VNS | 8.665 | 803 |
| **S14** | 20 Hz VNS | 9.353 | 996 |
| **S15** | 20 Hz VNS | 9.333 | 1000 |
| **S16** | 20 Hz VNS | 9.442 | 1000 |
| **S17** | 20 Hz VNS | 8.939 | 1000 |
| **S18** | 20 Hz VNS | 9.056 | 1000 |
| **S19** | 20 Hz VNS | 8.514 | 999 |
| **S20** | 20 Hz VNS | 8.844 | 1000 |
| **S21** | 30 Hz/ Standard VNS | 9.105 | 948 |
| **S22** | 30 Hz/ Standard VNS | 9.063 | 948 |
| **S23** | 30 Hz/ Standard VNS | 8.966 | 1000 |
| **S24** | 30 Hz/ Standard VNS | 8.702 | 1000 |
| **S25** | 30 Hz/ Standard VNS | 9.003 | 998 |
| **S26** | 30 Hz/ Standard VNS | 8.637 | 1000 |
| **S27** | 30 Hz/ Standard VNS | 8.848 | 1000 |
| **S28** | 30 Hz/ Standard VNS | 9.028 | 1000 |
| **S29** | 30 Hz/ Standard VNS | 9.186 | 1000 |
| **S30** | 30 Hz/ Standard VNS | 8.901 | 1000 |
| **S31** | 45 Hz VNS | 9.129 | 1000 |
| **S32** | 45 Hz VNS | 9.154 | 960 |
| **S33** | 45 Hz VNS | 8.954 | 1000 |
| **S34** | 45 Hz VNS | 9.640 | 997 |
| **S35** | 45 Hz VNS | 8.690 | 1000 |
| **S36** | 45 Hz VNS | 9.297 | 1000 |
| **S37** | 45 Hz VNS | 9.264 | 1000 |
| **S38** | 45 Hz VNS | 8.993 | 1000 |
| **S39** | 45 Hz VNS | 8.748 | 1000 |
| **S40** | 45 Hz VNS | 9.121 | 976 |
| **S41** | Burst VNS | 9.278 | 1000 |
| **S42** | Burst VNS | 8.712 | 1000 |
| **S43** | Burst VNS | 9.051 | 997 |
| **S44** | Burst VNS | 9.013 | 1000 |
| **S45** | Burst VNS | 8.714 | 1000 |
| **S46** | Burst VNS | 9.386 | 854 |
| **S47** | Burst VNS | 9.088 | 1000 |
| **S48** | Burst VNS | 8.509 | 1000 |
| **S49** | Burst VNS | 8.972 | 1000 |
| **S50** | Burst VNS | 8.738 | 1000 |
| **S51** | Dispersed VNS | 9.212 | 989 |
| **S52** | Dispersed VNS | 8.914 | 1000 |
| **S53** | Dispersed VNS | 8.900 | 1000 |
| **S54** | Dispersed VNS | 9.879 | 912 |
| **S55** | Dispersed VNS | 8.748 | 1000 |
| **S56** | Dispersed VNS | 8.812 | 1000 |
| **S57** | Dispersed VNS | 8.644 | 1000 |
| **S58** | Dispersed VNS | 8.865 | 999 |
| **S59** | Dispersed VNS | 9.247 | 988 |
| **S60** | Dispersed VNS | 9.439 | 907 |

During training sessions paired with VNS, the mean of the interval between stimulations for each session was recorded. A maximum of 1000 stimulations were delivered. Subjects in the Sham group did not receive VNS.

### Table S2. ICMS data for all subjects

| Subject | Group | Cortical area of movement representation ($mm^2$) | | | | | Mean ICMS Threshold (µA) |
|---|---|---|---|---|---|---|---|
| | | Forelimb | Jaw | Vibrissa | Neck | Hindlimb | |
| **S1** | Sham | 7.50 | 1.00 | 0.50 | 0.25 | 1.25 | 112.62 |
| **S2** | Sham | 6.75 | 1.75 | 0.50 | 0.00 | 1.75 | 127.21 |
| **S3** | Sham | 6.75 | 2.00 | 1.50 | 0.00 | 2.25 | 107.60 |
| **S4** | Sham | 6.00 | 1.50 | 2.50 | 0.25 | 1.50 | 167.23 |
| **S5** | Sham | 4.25 | 1.50 | 1.00 | 0.00 | 1.00 | 160.65 |
| **S6** | Sham | 8.00 | 0.50 | 0.25 | 0.00 | 1.75 | 89.29 |
| **S7** | Sham | 8.75 | 0.75 | 1.75 | 0.00 | 2.50 | 71.45 |
| **S8** | Sham | 8.00 | 1.75 | 1.50 | 0.50 | 2.00 | 128.18 |
| **S9** | Sham | 8.50 | 1.75 | 0.00 | 0.00 | 2.25 | 125.40 |
| **S10** | Sham | 5.75 | 1.75 | 1.00 | 0.00 | 2.75 | 150.22 |
| **S11** | 20 Hz VNS | 5.75 | 2.25 | 0.75 | 0.00 | 1.25 | 119.00 |
| **S12** | 20 Hz VNS | 7.50 | 1.00 | 0.75 | 0.00 | 1.00 | 92.44 |
| **S13** | 20 Hz VNS | 6.00 | 4.00 | 2.50 | 0.50 | 3.75 | 118.36 |
| **S14** | 20 Hz VNS | 5.25 | 2.50 | 0.25 | 0.25 | 2.25 | 141.19 |
| **S15** | 20 Hz VNS | 9.25 | 2.25 | 0.75 | 0.50 | 2.00 | 104.41 |
| **S16** | 20 Hz VNS | 6.75 | 1.75 | 0.75 | 0.50 | 0.25 | 107.00 |
| **S17** | 20 Hz VNS | 7.00 | 1.75 | 1.25 | 0.25 | 2.50 | 111.57 |
| **S18** | 20 Hz VNS | 7.00 | 3.00 | 0.00 | 0.00 | 1.00 | 106.14 |
| **S19** | 20 Hz VNS | 5.50 | 0.50 | 0.75 | 0.00 | 1.50 | 110.30 |
| **S20** | 20 Hz VNS | 7.50 | 2.75 | 1.50 | 0.00 | 3.25 | 88.60 |
| **S21** | 30 Hz/ Standard VNS | 8.25 | 1.25 | 0.25 | 0.00 | 1.25 | 91.82 |
| **S22** | 30 Hz/ Standard VNS | 7.25 | 2.00 | 0.25 | 0.00 | 1.25 | 194.65 |
| **S23** | 30 Hz/ Standard VNS | 8.50 | 3.00 | 1.25 | 0.00 | 3.75 | 95.91 |
| **S24** | 30 Hz/ Standard VNS | 8.00 | 3.00 | 1.50 | 0.25 | 1.50 | 97.89 |
| **S25** | 30 Hz/ Standard VNS | 6.00 | 4.00 | 2.25 | 0.25 | 3.00 | 135.81 |
| **S26** | 30 Hz/ Standard VNS | 6.25 | 3.50 | 1.00 | 0.00 | 3.50 | 134.04 |
| **S27** | 30 Hz/ Standard VNS | 7.50 | 0.50 | 0.75 | 0.00 | 0.75 | 113.95 |
| **S28** | 30 Hz/ Standard VNS | 6.50 | 2.25 | 1.00 | 0.00 | 2.50 | 82.65 |
| **S29** | 30 Hz/ Standard VNS | 7.50 | 1.75 | 1.25 | 0.00 | 2.25 | 94.12 |
| **S30** | 30 Hz/ Standard VNS | 6.75 | 3.50 | 1.75 | 0.00 | 1.25 | 143.77 |
| **S31** | 45 Hz VNS | 6.00 | 1.25 | 1.25 | 0.25 | 0.75 | 102.37 |
| **S32** | 45 Hz VNS | 8.75 | 3.00 | 0.50 | 0.00 | 3.50 | 125.87 |
| **S33** | 45 Hz VNS | 8.75 | 1.75 | 3.25 | 0.00 | 1.75 | 73.06 |
| **S34** | 45 Hz VNS | 5.25 | 2.25 | 1.25 | 0.00 | 0.75 | 178.16 |
| **S35** | 45 Hz VNS | 7.75 | 1.25 | 0.50 | 0.25 | 1.75 | 131.52 |
| **S36** | 45 Hz VNS | 6.25 | 4.00 | 0.50 | 0.50 | 1.75 | 146.35 |
| **S37** | 45 Hz VNS | 7.00 | 3.50 | 1.75 | 0.00 | 3.00 | 117.87 |
| **S38** | 45 Hz VNS | 6.00 | 1.25 | 0.75 | 0.00 | 2.25 | 110.24 |
| **S39** | 45 Hz VNS | 5.75 | 2.50 | 1.00 | 0.00 | 2.00 | 116.00 |
| **S40** | 45 Hz VNS | 7.50 | 0.75 | 0.25 | 0.00 | 1.50 | 101.50 |
| **S41** | Burst VNS | 6.75 | 3.00 | 1.00 | 0.00 | 1.75 | 151.20 |
| **S42** | Burst VNS | 7.25 | 2.00 | 1.75 | 0.75 | 2.00 | 121.64 |
| **S43** | Burst VNS | 4.50 | 0.75 | 0.00 | 0.00 | 0.25 | 135.91 |
| **S44** | Burst VNS | 8.25 | 1.25 | 0.50 | 0.00 | 2.00 | 93.54 |
| **S45** | Burst VNS | 6.25 | 2.00 | 0.00 | 0.00 | 2.00 | 113.17 |
| **S46** | Burst VNS | 6.00 | 1.50 | 1.00 | 0.25 | 2.75 | 101.09 |
| **S47** | Burst VNS | 7.25 | 0.75 | 0.25 | 0.00 | 1.75 | 185.25 |
| **S48** | Burst VNS | 10.25 | 1.25 | 0.75 | 0.00 | 1.75 | 91.79 |
| **S49** | Burst VNS | 8.50 | 1.75 | 1.25 | 0.00 | 2.75 | 99.02 |
| **S50** | Burst VNS | 8.75 | 2.50 | 0.00 | 0.00 | 3.00 | 140.53 |
| **S51** | Dispersed VNS | 7.50 | 1.25 | 0.25 | 0.00 | 0.50 | 137.11 |
| **S52** | Dispersed VNS | 5.50 | 1.50 | 2.00 | 0.25 | 1.00 | 93.41 |
| **S53** | Dispersed VNS | 8.00 | 0.50 | 0.50 | 0.00 | 1.50 | 108.57 |
| **S54** | Dispersed VNS | 5.50 | 3.50 | 1.00 | 0.25 | 3.25 | 133.15 |
| **S55** | Dispersed VNS | 6.75 | 2.75 | 1.25 | 0.00 | 2.75 | 117.78 |
| **S56** | Dispersed VNS | 9.00 | 1.50 | 1.00 | 0.00 | 3.00 | 134.66 |
| **S57** | Dispersed VNS | 8.00 | 2.50 | 0.25 | 0.25 | 1.75 | 80.59 |
| **S58** | Dispersed VNS | 8.50 | 1.50 | 0.75 | 0.25 | 2.00 | 150.00 |
| **S59** | Dispersed VNS | 6.50 | 1.75 | 0.50 | 0.00 | 2.50 | 105.78 |
| **S60** | Dispersed VNS | 8.75 | 1.75 | 0.00 | 0.25 | 1.50 | 97.96 |

Each row represents an individual subject. ICMS elicited movements in forelimb, jaw, vibrissa, neck, and hindlimb in areas of the specified size. Mean ICMS threshold is the mean of the minimum currents required to produce movement at each stimulation site across the map.

**A**

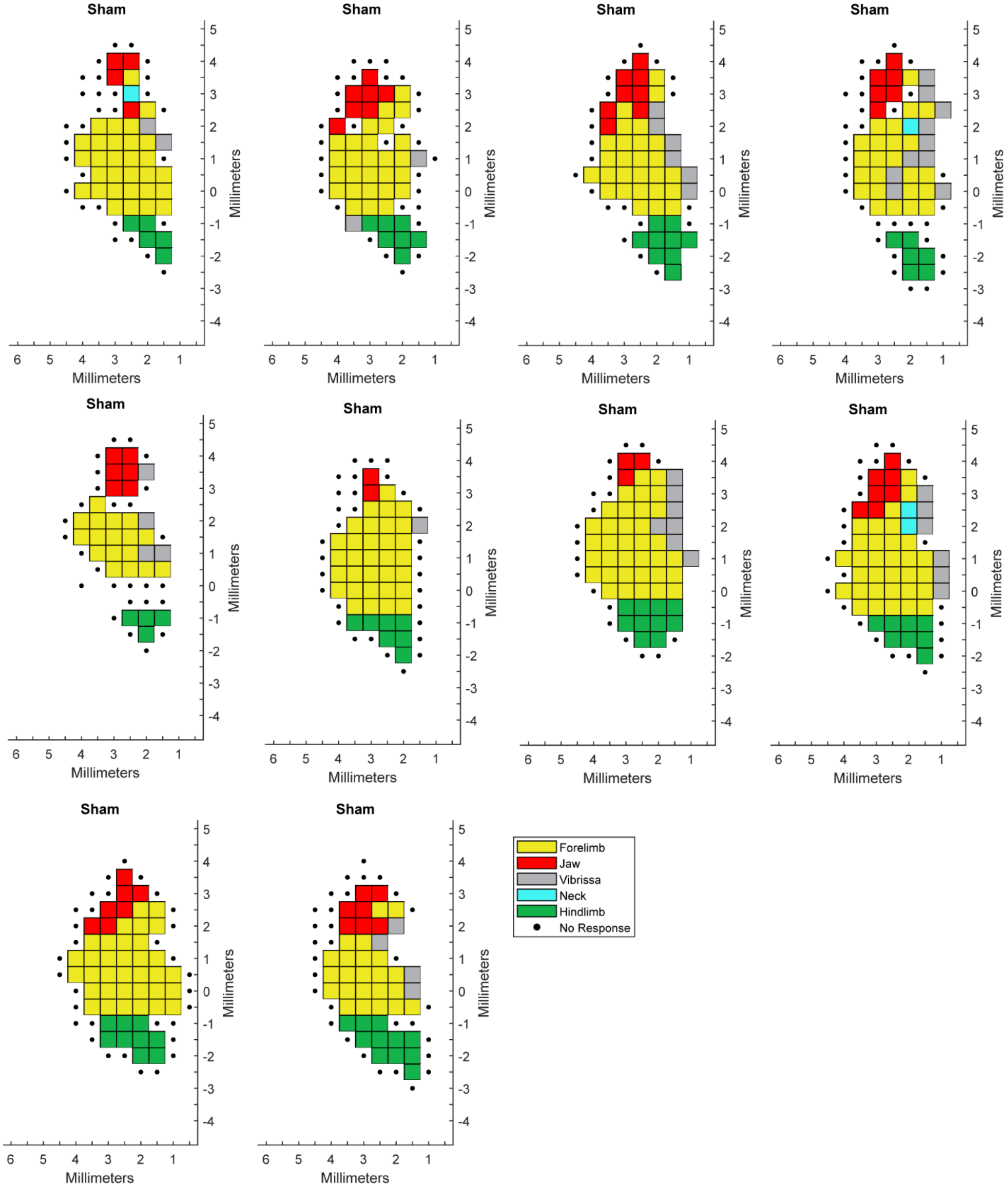
Sham
Millimeters
Forelimb
Jaw
Vibrissa
Neck
Hindlimb
No Response

**B**

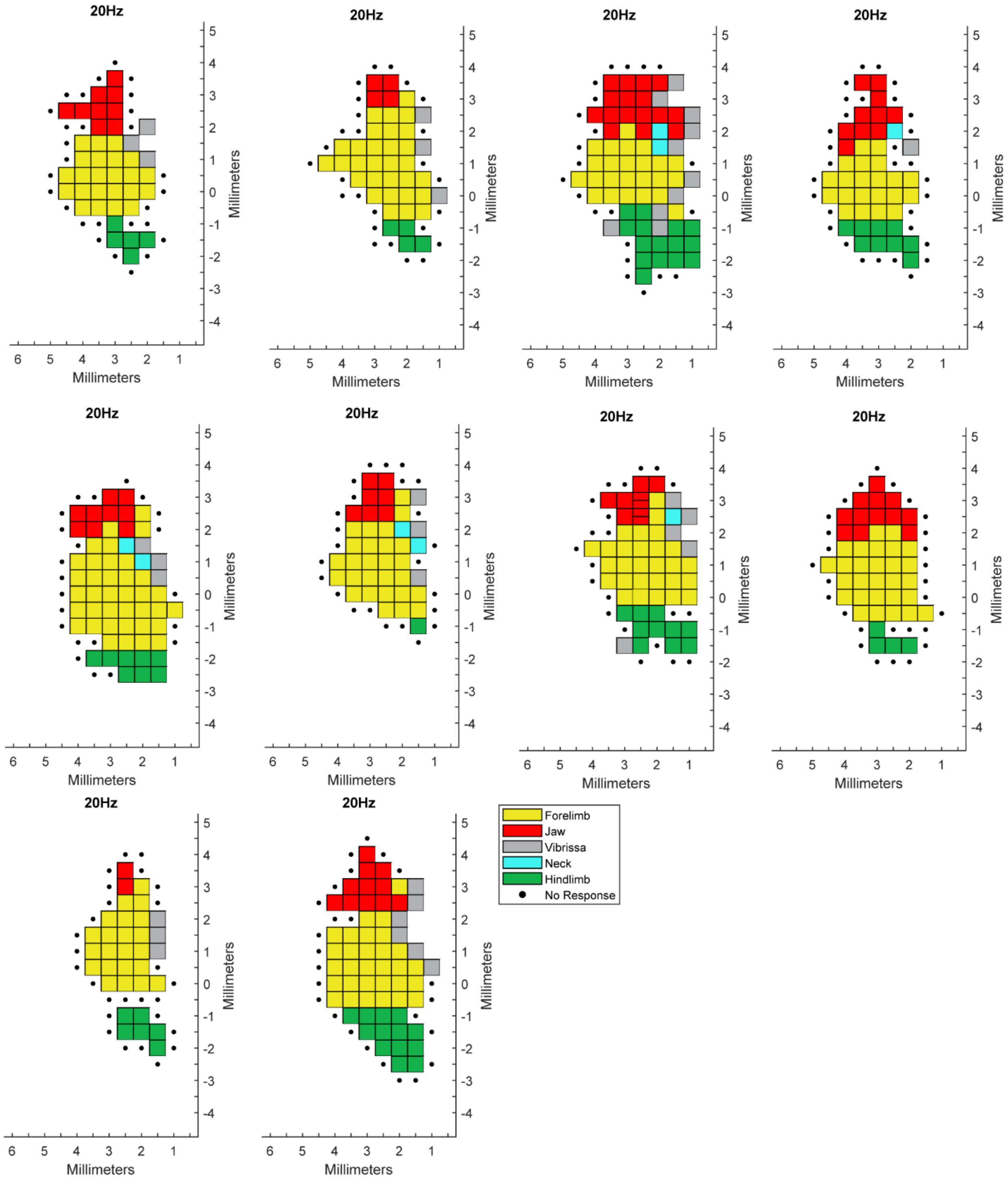
20Hz
Millimeters
Forelimb
Jaw
Vibrissa
Neck
Hindlimb
No Response

**C**

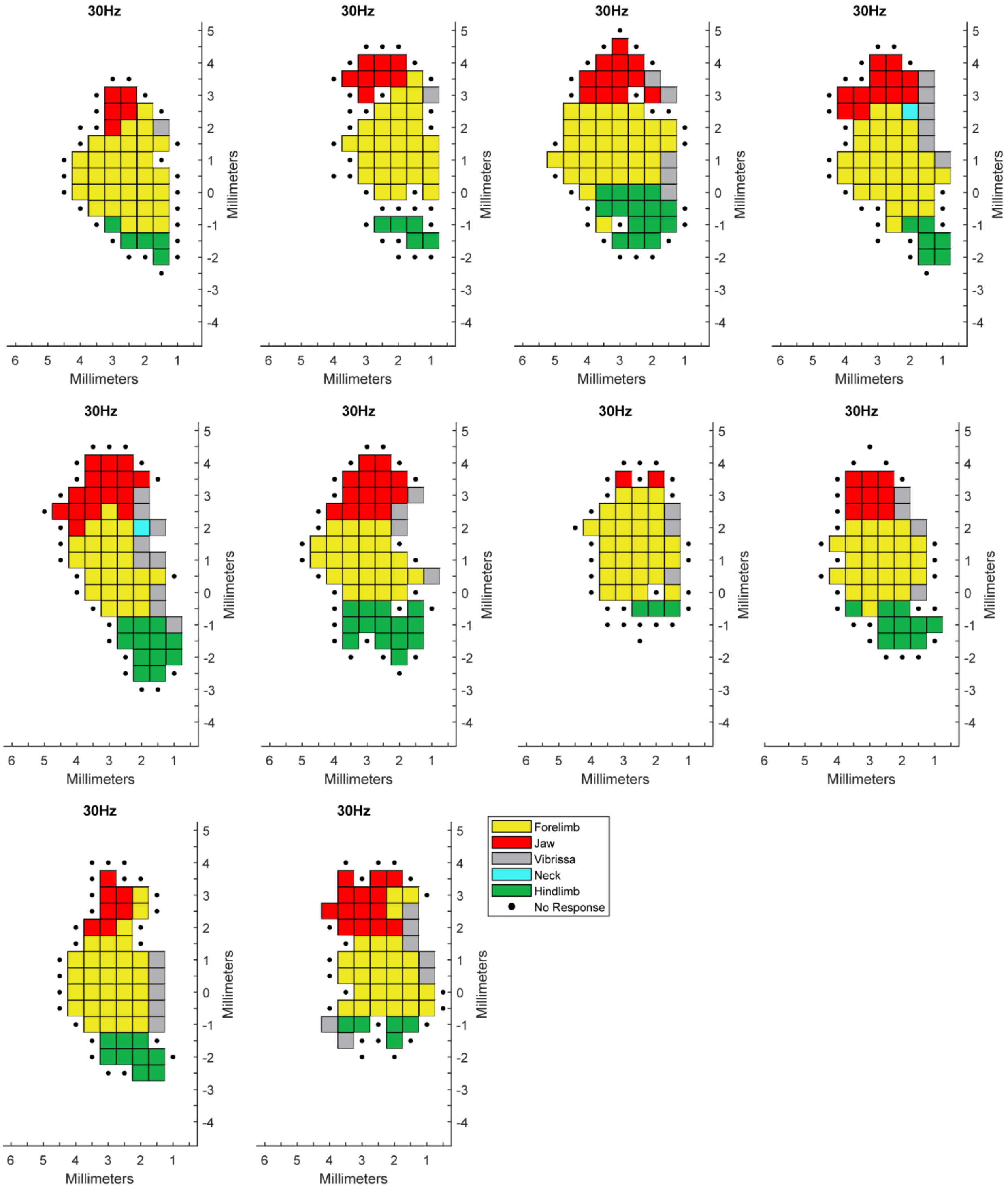
30Hz
Millimeters
Millimeters
Forelimb
Jaw
Vibrissa
Neck
Hindlimb
No Response

D

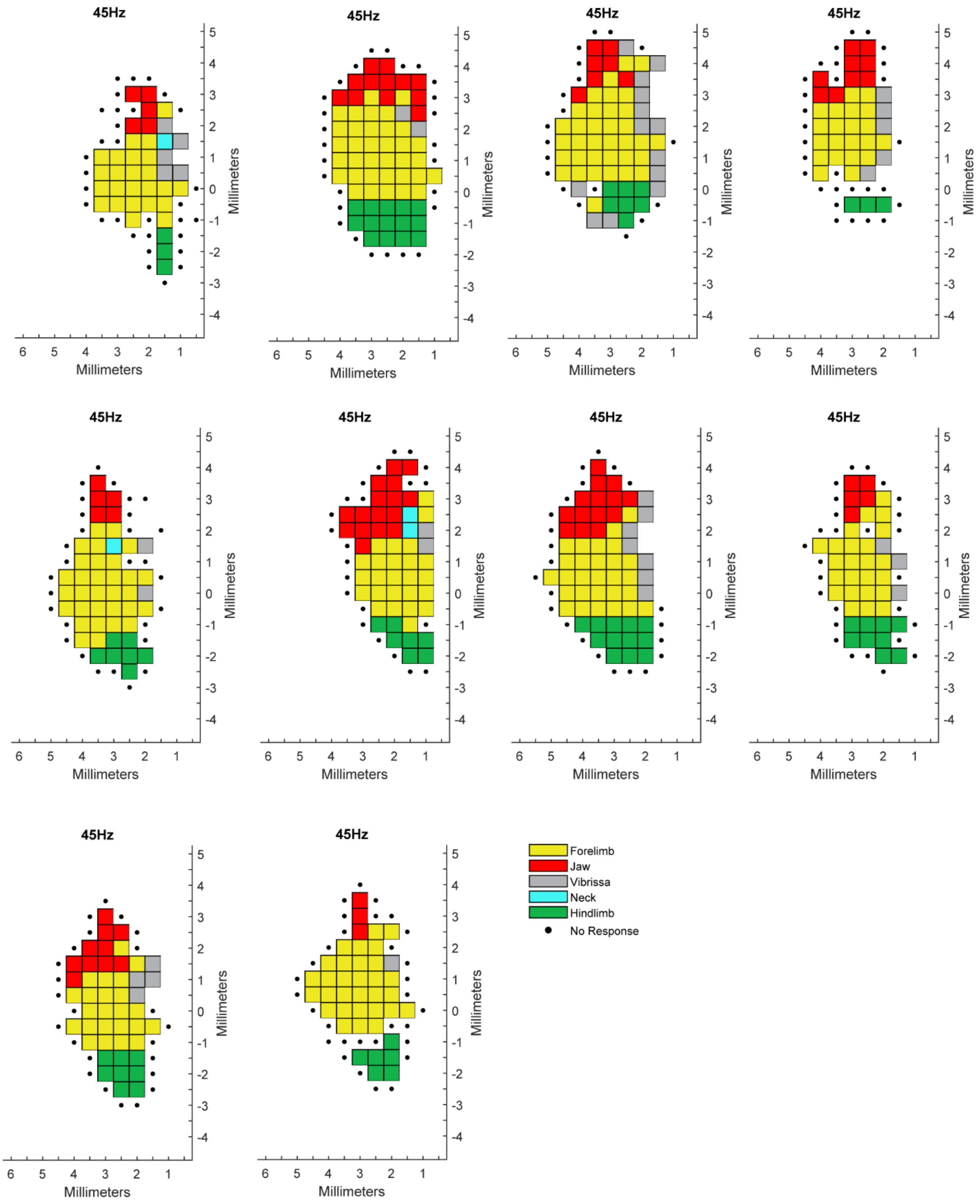
45Hz
Millimeters
Millimeters
Forelimb
Jaw
Vibrissa
Neck
Hindlimb
No Response

**E**

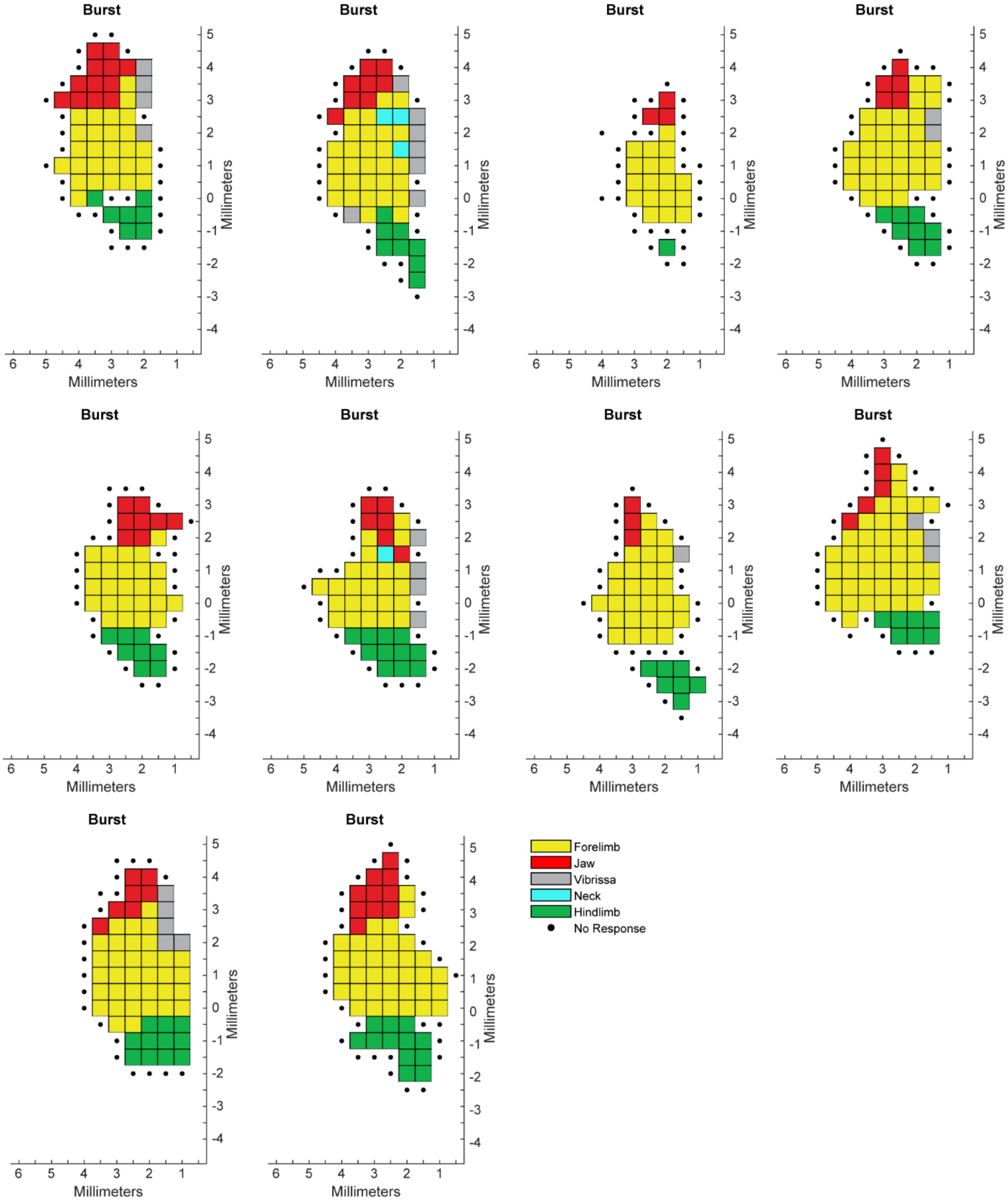

Burst
Burst
Burst
Burst
Burst
Burst
Burst
Burst
Burst
Burst
Millimeters
Millimeters
5 4 3 2 1 0 -1 -2 -3 -4
6 5 4 3 2 1
Forelimb
Jaw
Vibrissa
Neck
Hindlimb
No Response

F

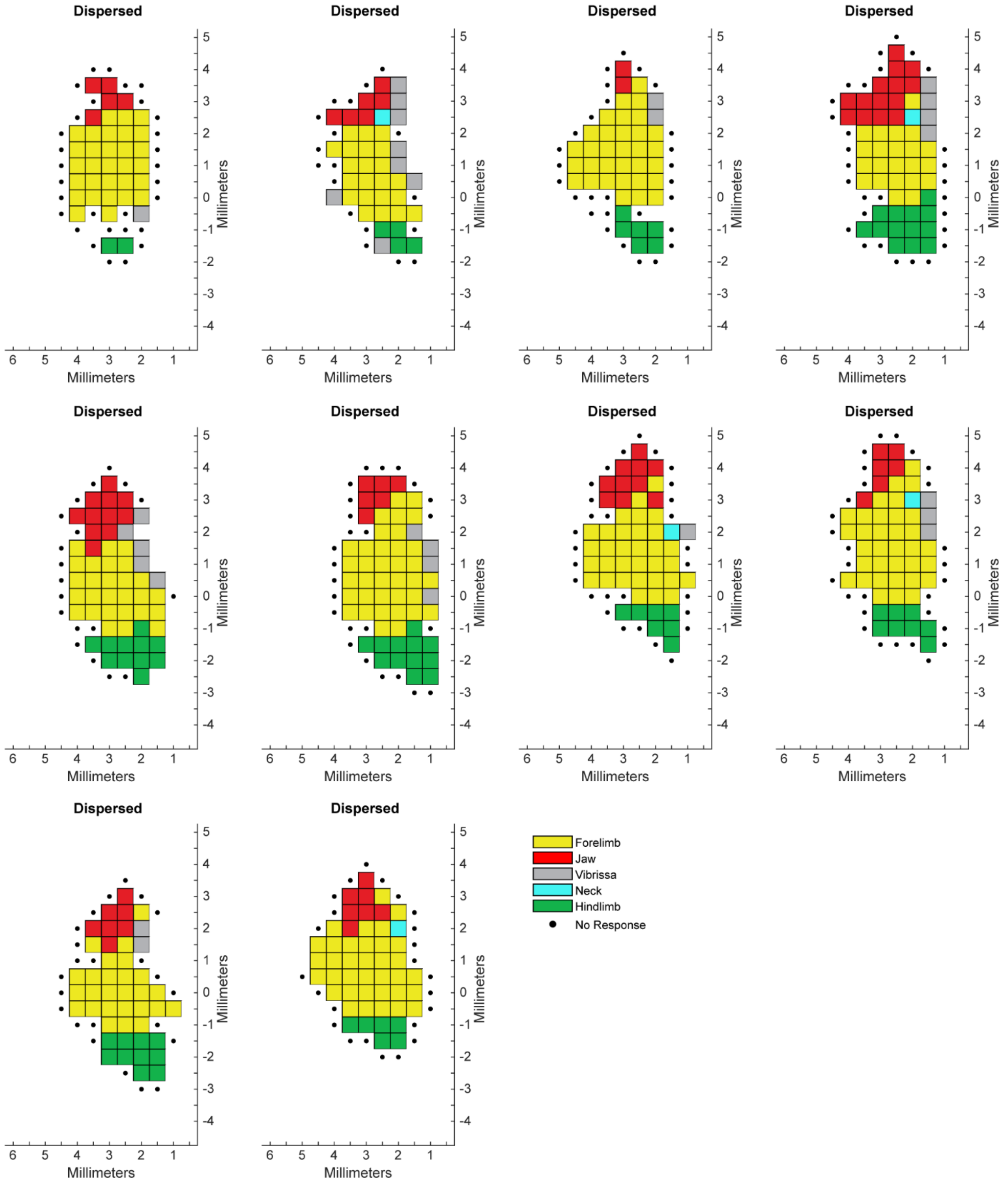


**Figure S1. Raw ICMS maps for all subjects.** Each subpanel illustrates the ICMS map from a single subject. Groups are presented in different panels: (A) Sham (B) 20 Hz VNS, (C) 30 Hz (Standard) VNS, (D) 45 Hz VNS, (E) Burst VNS, and (F) Dispersed VNS. On each subpanel, axes denote stereotaxic coordinates relative to bregma, with rostro-caudal directions from +5mm to -4mm and mediolateral directions from 0mm to +6mm.

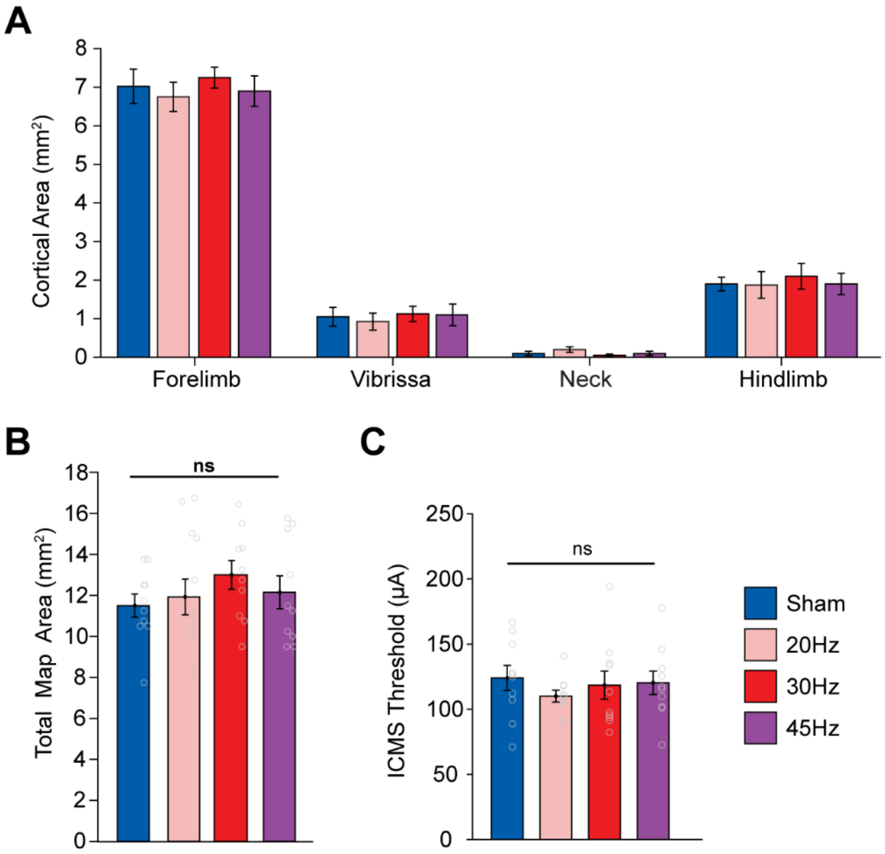


**Figure S2. Other movement representations and ICMS thresholds are not changed by moderate VNS frequencies.** (A) Other (non-jaw) cortical movement representations were not affected by moderate frequencies of VNS. (B) Additionally, moderate VNS frequencies did not change the total map area (B) or ICMS thresholds (C). Gray circles depict individual subjects. Bars represent mean ± SEM. * denotes $p < 0.0167$; n.s., not significant.

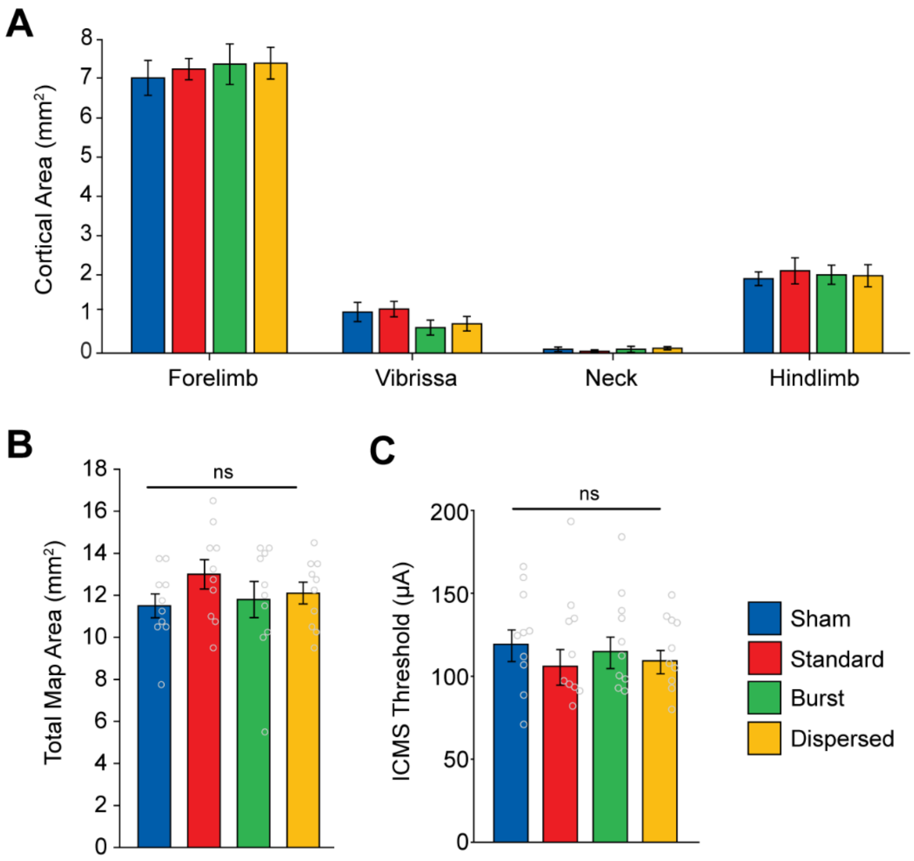


**Figure S3. Other movement representations and ICMS thresholds are not changed by burst and dispersed VNS paradigms.** (A) Other (non-jaw) cortical movement representations were not affected by different temporal VNS paradigms. (B) Additionally, burst and dispersed VNS did not change the total map area (B) or ICMS thresholds (C). Gray circles depict individual subjects. Bars represent mean ± SEM. * denotes $p < 0.0167$; n.s., not significant.
